## Supplementary figures and tables for "Investigating the interactions of the cucumber mosaic virus 2b protein with the viral 1a replicase component and the cellular RNA silencing factor Argonaute 1"

**Table S1. Primer sequences.**

| Primer name | Sequence (5'-3' ) |
|---|---|
| Gateway cloning | |
| 2b attB Rv | GGGGACCACTTTGTACAAGAAAGCTGGGTTGAAAGCACCTTC CGCCCA |
| 2b attB Fw | GGGGACAAGTTTGTACAAAAAAGCAGGCTTCATGGAATTG AACGTAGGT |
| 2bD1-17 Fw | GGGGACAAGTTTGTACAAAAAAGCAGGCTTCATGGTGGA GGCGAAGAAGC |
| 2bΔ95-110 Rv | GGGGACCACTTTGTACAAGAAAGCTGGGTTAAAATCATG GTCTTCCGCCGA |
| 2bΔ85-110 | GGGGACCACTTTGTACAAGAAAGCTGGGTTACGAGAGGCCTC AGACTCG |
| 2bΔ74-110 | GGGGACCACTTTGTACAAGAAAGCTGGGTTCACATGGCGGCA TGAC |
| 2bΔ56-110 | GGGGACCACTTTGTACAAGAAAGCTGGGTTTAGGAAGCGGAA TAGTCTGAGATTTGAAC |
| 2bΔ83-110 Rv | GGGGACCACTTTGTACAAGAAAGCTGGGTTTCAGGCCTCAGA CTCGGG |
| 2bΔ1-55 Fw | GGGGACAAGTTTGTACAAAAAAGCAGGCTTAATGCCGTTCTA TCAAGTGGATGGTTCG |
| 2bΔ61-110 Rv | GGGGACCACTTTGTACAAGAAAGCTGGGTTCACTTGATAGAA CGGTAGGAAGCG |
| 2bΔ65-110 Rv | GGGGACCACTTTGTACAAGAAAGCTGGGTTTTCCGAACCATC CACTTGATAGAACG |
| 2bΔ69-110 Rv | GGGGACCACTTTGTACAAGAAAGCTGGGTTTGACCCTGTCAG TTCCGAACC |
| Q5 mutagenesis | |
| 2bLS/Fny83-93 Rv | GGCCTCAGACTCGGGTAA |
| 2bLS/Fny83-93 Fw | TTTGACGATACAGATTGGTTCG |
| 2bΔ56-60/65 Rv | TAGGAAGCGGAATAGTCTGAG |
| 2bΔ39-48 Fw | CTCAGACTATTCCGCTTCCTAC |
| 2bΔ39-48 Rv | GTGACCTCGTTCCCGTCG |
| 2bΔ56-65 Fw | ACAGGGTCATGCCGC |
| 2bΔ56-60 Fw | GATGGTTCGGAACTGACAG |
| 2b56aaa65 Rv | CGCCGCCGCCGCCGCCGCCGCCGCCGCCGCTAGGAAGCGGAA TAGTCTGAG |
| 2b56aaa60 Rv | CGCCGCCGCCGCCGCTAGGAAGCGGAATAGTCTGAG |
| 2bFny/LS56-60 Fw | CCGTTCcatggAGTGGATGGTTCGGAA |
| 2bΔ83-93 (step 2) Rv | GAAAGCACCTTCCGCCCATTCGTTACCGGCGAACCAATCTGT ATCGTCAAAGGCCTC |
| 2bΔ83-93 (step 1) Rv | CCAATCTGTATCGTCAAAGGCCTCAGACTCGGGTAAC |
| 2b Fw | ATGGAATTGAACGTAGGTGCAATGAC |

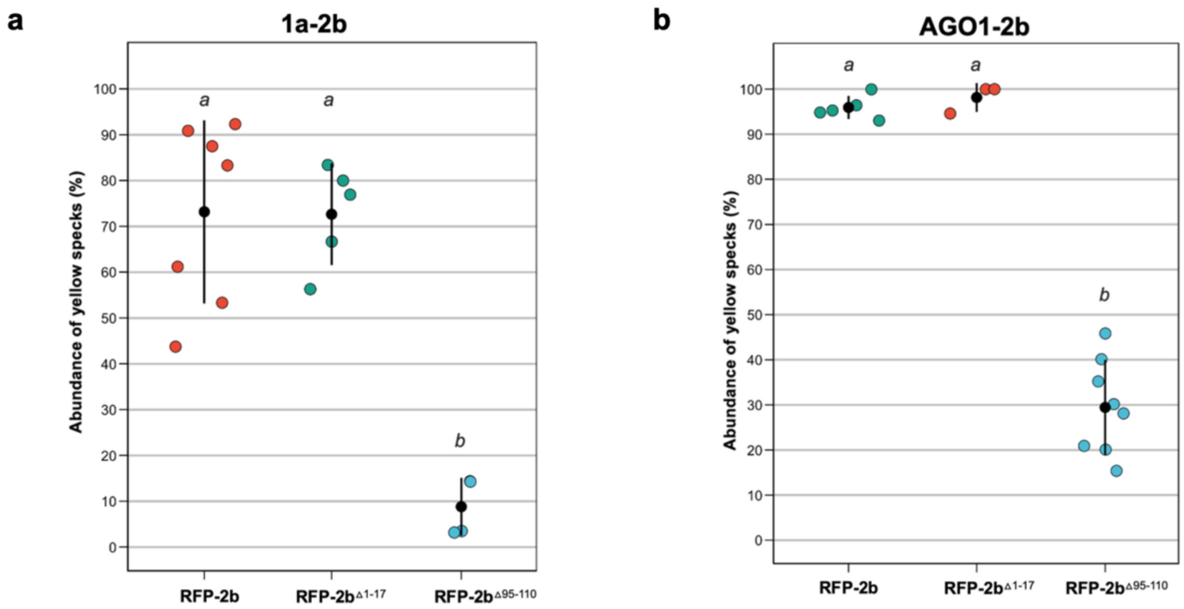


**Fig. S1** The C-terminal domain of the cucumber mosaic virus 2b protein enhances its interaction with the Fny-CMV 1a and AGO1 proteins. Red fluorescent protein (RFP)-tagged CMV 2b proteins containing deletions in their N-terminal domain (RFP-2b$^{\Delta 1\text{-}17}$) or C-terminal domain (RFP-2b$^{\Delta 95\text{-}110}$) were transiently co-expressed with green fluorescent protein (GFP)-tagged Fny-CMV 1a or AGO1 proteins in agroinfiltrated leaves of *N. benthamiana*. The abundance of yellow specks in each image was quantified by calculating the number of yellow specks as a percentage of the total specks present in the image. Individual abundance values are presented as jitter plots with each mean value and standard error depicted as black bars. Since yellow specks result from the overlay of GFP and RFP fluorescent signals they were used as a measure of co-localisation between co-infiltrated GFP- and RFP-tagged proteins. **a** Co-localisation results for 1a-GFP with RFP-2b$^{\Delta 95\text{-}110}$, RFP-2b$^{\Delta 1\text{-}17}$ and RFP-2b. Lower case letters *a* and *b* indicate mean values for abundance of yellow specks that are significantly different from each other (P <0.0001: Tukey's multiple comparison of means). Values labelled with the same letter are not significantly different form each other. Deletion of the N-terminal domain of the 2b protein had no significant effect on colocalisation with CMV 1a proteins. However, deletion of the C-terminal domain of the 2b protein significantly reduced colocalisation with CMV 1a proteins. **b** Co-localisation results for AGO1-GFP with RFP-2b$^{\Delta 95\text{-}110}$, RFP-2b$^{\Delta 1\text{-}17}$ and RFP- 2b. Lower case letters *a* and *b* indicate mean values for abundance of yellow specks that are significantly different from each other (P <0.0001: Tukey's multiple comparison of means). Values labelled with the same letter are not significantly different form each other. Deletion of the N-terminal domain of 2b had no significant effect on colocalisation with AGO1 proteins. However, deletion of the C-terminal domain of 2b significantly reduced colocalisation with AGO1 proteins. Number of independent leaves imaged for each treatment, $n \geq 3$.

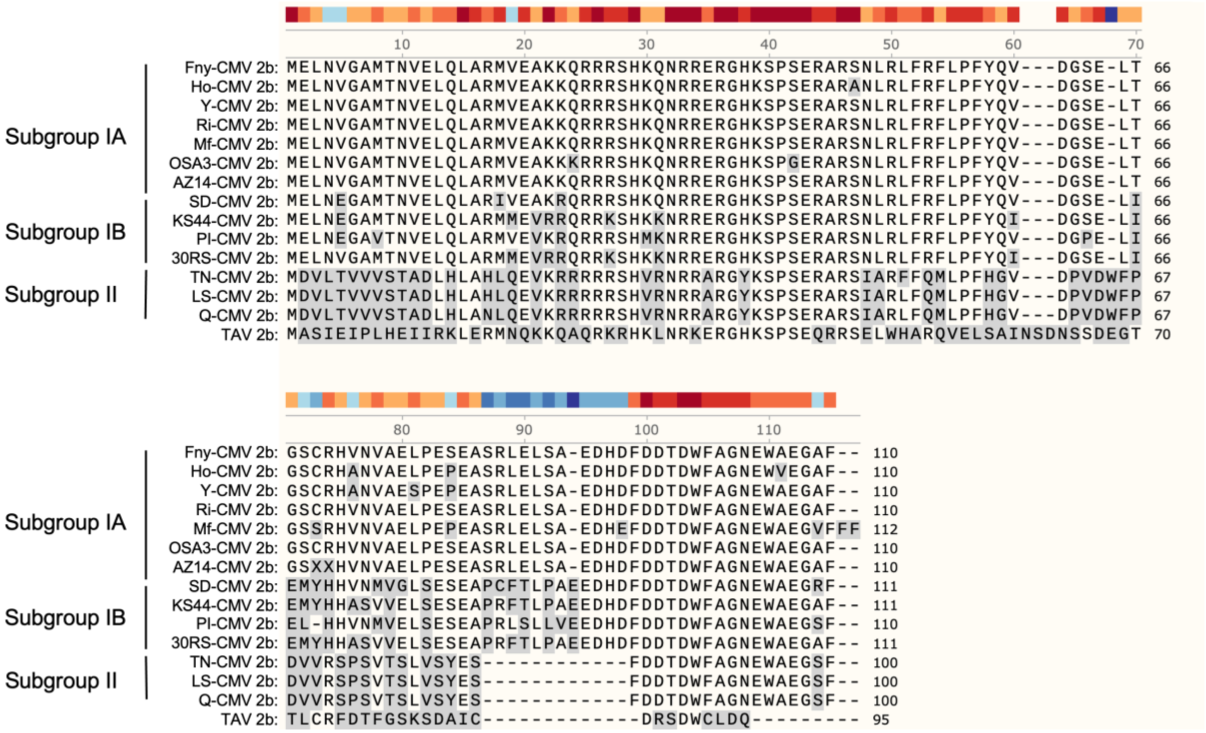


**Fig. S2** The amino acid sequences of 2b proteins encoded by a selection of strains of *Cucumber mosaic virus* (CMV) from Subgroups IA, IB and II and the 2b protein from *Tomato aspermy virus* (TAV). The Fny-CMV 2b protein sequence is given as a reference to compare with 2b protein sequences from a small number of other CMV strains to highlight certain differences and similarities between 2b proteins of viruses in Subgroups IA, IB and II. Differences in 2b protein primary sequences between the Fny-CMV and other strains are highlighted in grey. The numbering of amino acid residues is based on the Fny-CMV 2b protein sequence. The GenBank accession numbers for the sequences used in this alignment are NC002035 for Fny-CMV (2b-Fny), D12538 for Y-CMV (2b-Y), LC593245 for Ho- CMV (2b- Ho), AM183118 for RI-8-CMV (2b-RI-8), AJ276480 for Mf-CMV (2b-Mf), HE971489 for OSA3-CMV (2b-OSA3), QBH72281 for AZ14-CMV (2b-AZ14), D86330 for SD-CMV, CBG76802 for KS44-CMV (2b-KS44), CAJ65577 for PI-1-CMV (2b-PI-1), FN552601 for 30RS-CMV (2b-30RS), BAD15371 for TN-CMV (2b-TN), AF416900 for LS-CMV (2b- LS), Q66125 for Q-CMV (2b-Q) and AJ320274 for KC-TAV (TAV 2b).

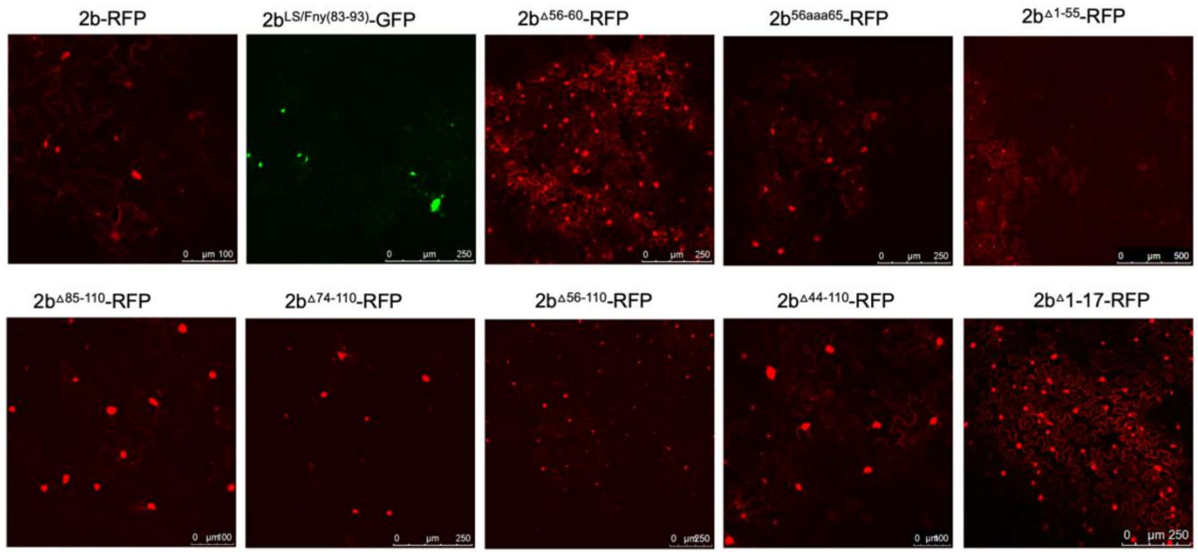


**Fig. S3** Mutations affecting the subcellular localization of cucumber mosaic virus 2b proteins. Agroinfiltration was used for transient expression in *N. benthamiana* leaves of C-terminal red fluorescent protein 2b-(RFP) or C-terminal green fluorescent protein 2b-(GFP) fusion protein. All mutant versions of the 2b protein showed nuclear and cytoplasmic localisation. However, mutant versions of the 2b protein lacking residues at their C-terminus (such as 85-110, 74-110, 56-110 or 44-110) showed an increased nuclear localisation compared to the wild-type protein. Increased nuclear localisation was also seen for the LS/Fny chimeric mutant in which residues present in the Fny-CMV 2b protein (residues 83-93) but absent in the sequence of the LS-CMV 2b protein, were introduced into a chimeric LS-CMV protein. However, the N-terminal mutant from residues 1-17, mutations from 56-60 or substitution of alanine from 56-60 had no impact on the localisation of CMV 2b. In contrast, deletion of residues from 1-55 caused a reduction in nuclear localisation and an increase in cytoplasmic localisation.

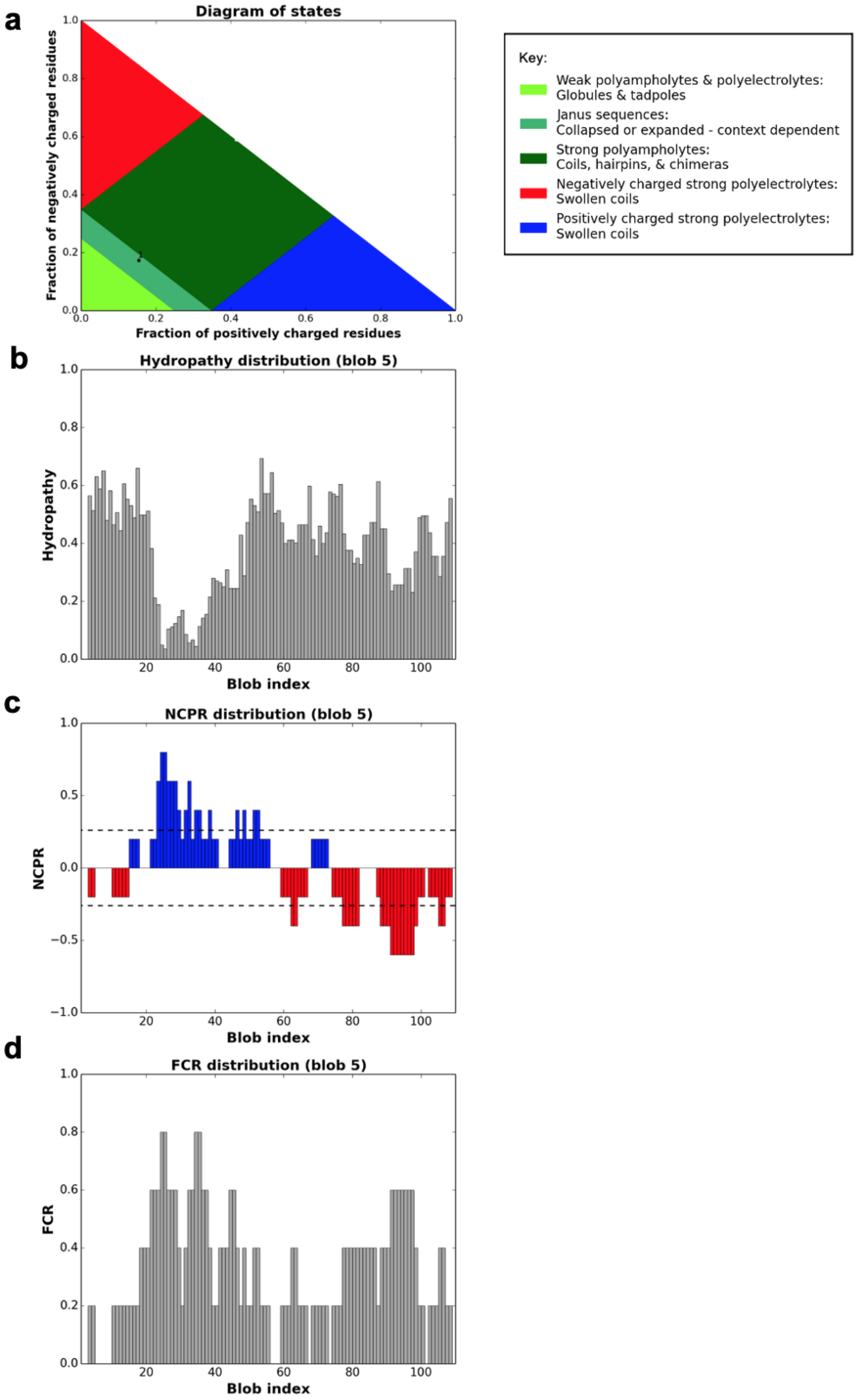
a
Diagram of states
Fraction of negatively charged residues
Fraction of positively charged residues
Key:
Weak polyampholytes & polyelectrolytes:
Globules & tadpoles
Janus sequences:
Collapsed or expanded - context dependent
Strong polyampholytes:
Coils, hairpins, & chimeras
Negatively charged strong polyelectrolytes:
Swollen coils
Positively charged strong polyelectrolytes:
Swollen coils
b
Hydropathy distribution (blob 5)
Hydropathy
Blob index
c
NCPR distribution (blob 5)
NCPR
Blob index
d
FCR distribution (blob 5)
FCR
Blob index

**Fig. S4** Prediction of physicochemical properties for the Fny-CMV 2b protein. This figure shows the predictions of the CIDER server (http://pappulab.wustl.edu/CIDER/about /). **a** The diagram of states provides an estimate of the kinds of ensembles adopted by an intrinsically disordered sequence. The predictions suggest that the Fny-CMV 2b protein sequence has a context-dependent structure, existing in either collapsed or expanded states depending on other factors such as salt concentration, ligand binding, *cis*- interactions etc. **b** The mean hydropathy of the sequence shown as a re-scaled Kyte-Doolittle (Kyte and Doolittle, 1982) hydropathy value which lies between 0 (least hydrophobic) and 1 (most hydrophobic). **c** The net charge per residue (NCPR) predicts that the Fny- CMV 2b protein has clearly grouped regions of positive and negative charge. **d** The fraction of charged residues in the sequence (FCR) also shows grouping of regions with charged residues. This may facilitate self-interaction and the regions of positive charge are likely candidates for RNA interaction.

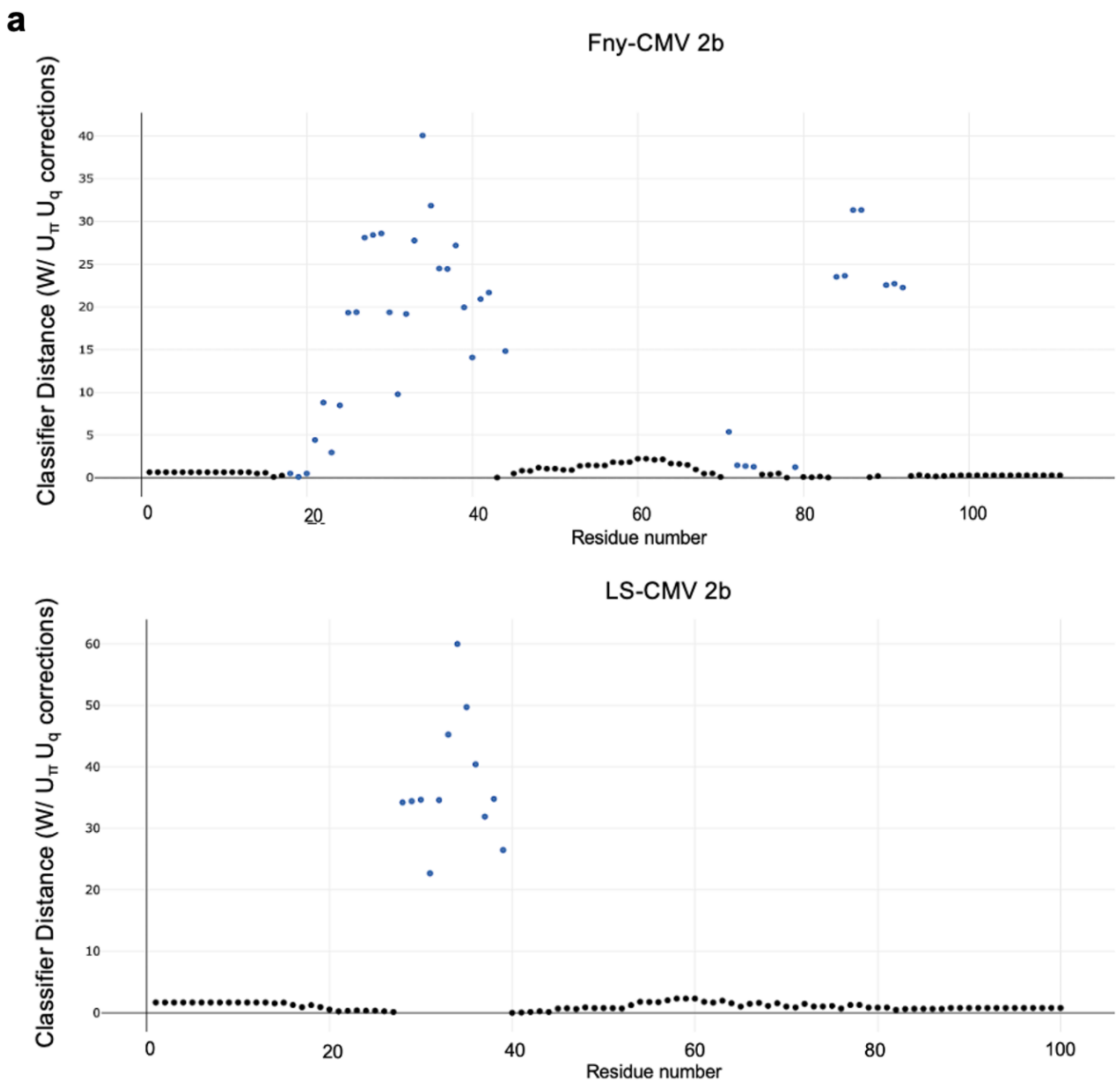
a
Fny-CMV 2b
Classifier Distance (W/ $U_\pi$ $U_q$ corrections)
40
35
30
25
20
15
10
5
0
0
20
40
60
80
100
Residue number
LS-CMV 2b
Classifier Distance (W/ $U_\pi$ $U_q$ corrections)
60
50
40
30
20
10
0
0
20
40
60
80
100
Residue number

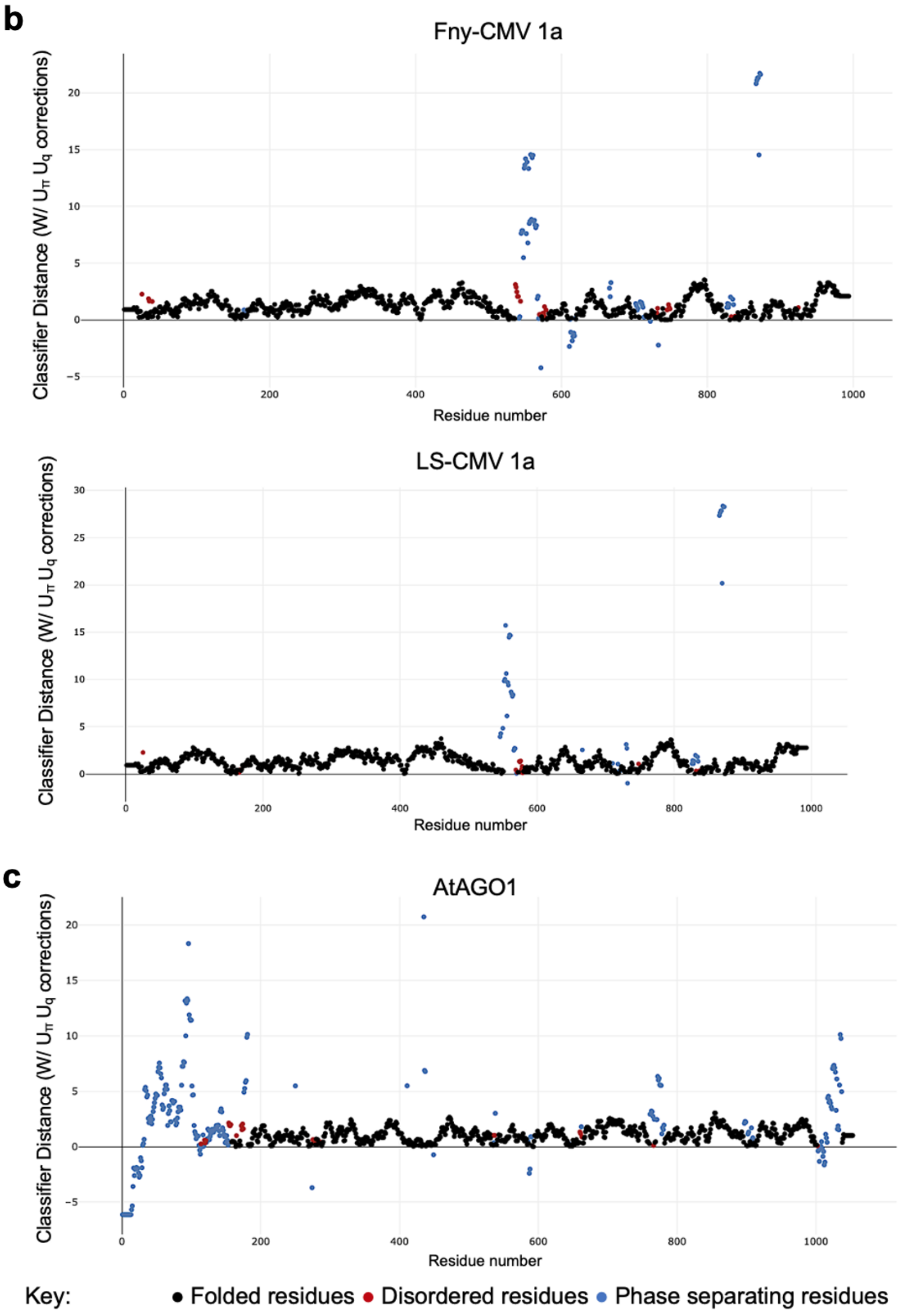
b
Fny-CMV 1a
Classifier Distance (W/ $U_\pi$ $U_q$ corrections)
Residue number
LS-CMV 1a
Classifier Distance (W/ $U_\pi$ $U_q$ corrections)
Residue number
c
AtAGO1
Classifier Distance (W/ $U_\pi$ $U_q$ corrections)
Residue number
Key:
Folded residues
Disordered residues
Phase separating residues

**Fig. S5** Predictions of phase separation. The ParSe v2 program identifies protein regions likely to exhibit physiological phase separation behaviour. The results shown here are for the modified ParSe v2 program accounting for the effects of interactions between amino acids (termed $U_\pi$ for π–π and cation–π interactions and $U_q$ for charge-based effects). Residues are colour coded based on their predicted state with black representing folded residues, red representing disordered residues and blue representing phase separating residues. The classifier distance on the Y-axis represents the confidence in the assigned states with higher values relating to higher confidence. **a** The Fny-CMV 2b protein is predicted to contain two regions which are promote phase separation behaviour while the LS-CMV 2b protein contains only one such region. **b** The Fny-CMV 1a protein contains residues which are likely to promote phase separating behaviour, but these represent a smaller percentage of residues than those seen in the Fny-CMV 2b protein. The LS-CMV 1a protein also contains residues which are likely to promote phase separation, but there are slightly fewer such residues than in the Fny-CMV 1a protein (41 compared with 66). **C,** The *A. thaliana* AGO1 protein is also predicted to contain residues promoting phase separation with the greatest density at its N-terminus.

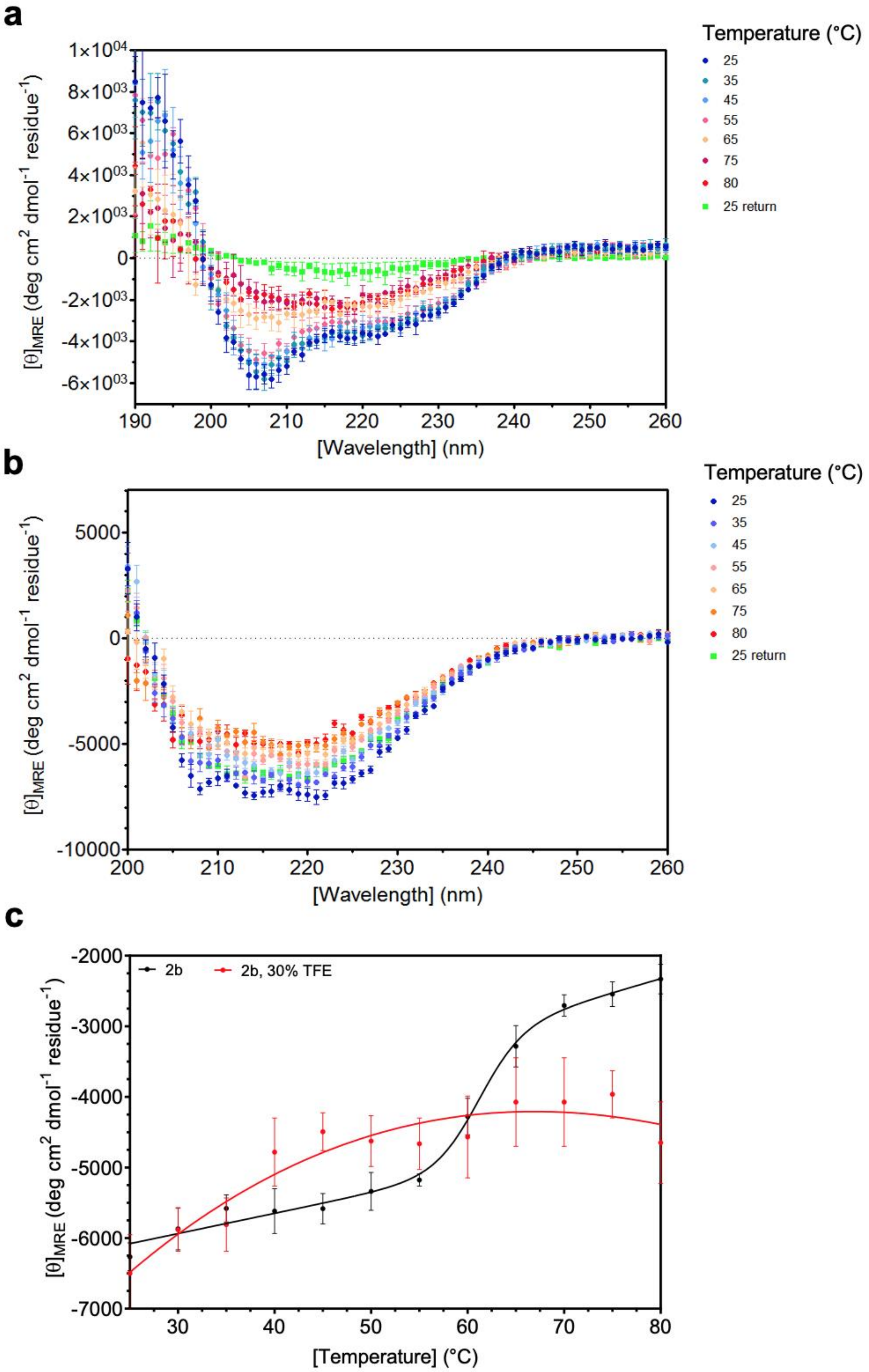
a
Temperature (°C)
25
35
45
55
65
75
80
25 return
$[\theta]_{MRE}$ (deg cm$^2$ dmol$^{-1}$ residue$^{-1}$)
1×10$^{04}$
8×10$^{03}$
6×10$^{03}$
4×10$^{03}$
2×10$^{03}$
0
-2×10$^{03}$
-4×10$^{03}$
-6×10$^{03}$
190
200
210
220
230
240
250
260
[Wavelength] (nm)
b
Temperature (°C)
25
35
45
55
65
75
80
25 return
$[\theta]_{MRE}$ (deg cm$^2$ dmol$^{-1}$ residue$^{-1}$)
5000
0
-5000
-10000
200
210
220
230
240
250
260
[Wavelength] (nm)
c
2b
2b, 30% TFE
$[\theta]_{MRE}$ (deg cm$^2$ dmol$^{-1}$ residue$^{-1}$)
-2000
-3000
-4000
-5000
-6000
-7000
30
40
50
60
70
80
[Temperature] (°C)

**Fig. S6** Temperature Denaturation of the Fny-CMV 2b protein. Circular dichroism data were obtained at a protein concentration of 13.5 µM 2b in 10 mM Phosphate buffer pH 7.5 at temperatures ranging from 25 °C to 80 °C. **a** Following heating, the 2b protein is irreversibly denatured as cooling (green symbols) does not revert the protein to its initial state at 25 °C (dark blue symbols). **b** Following the addition of 30% trifluroethanol (alpha-helical structure in proteins) the 2b protein does not completely unfold even when heated to 80 °C and following cooling back to 25 °C the protein does revert back to its initial folded state. **c** There is a transition between folded and unfolded state where 50% of the 2b protein is in each state at 61 °C when fitted to a Boltzmann model with sloping baselines (black symbols) measured at 208 nm. All results are shown as the mean of five scans ± SEM.

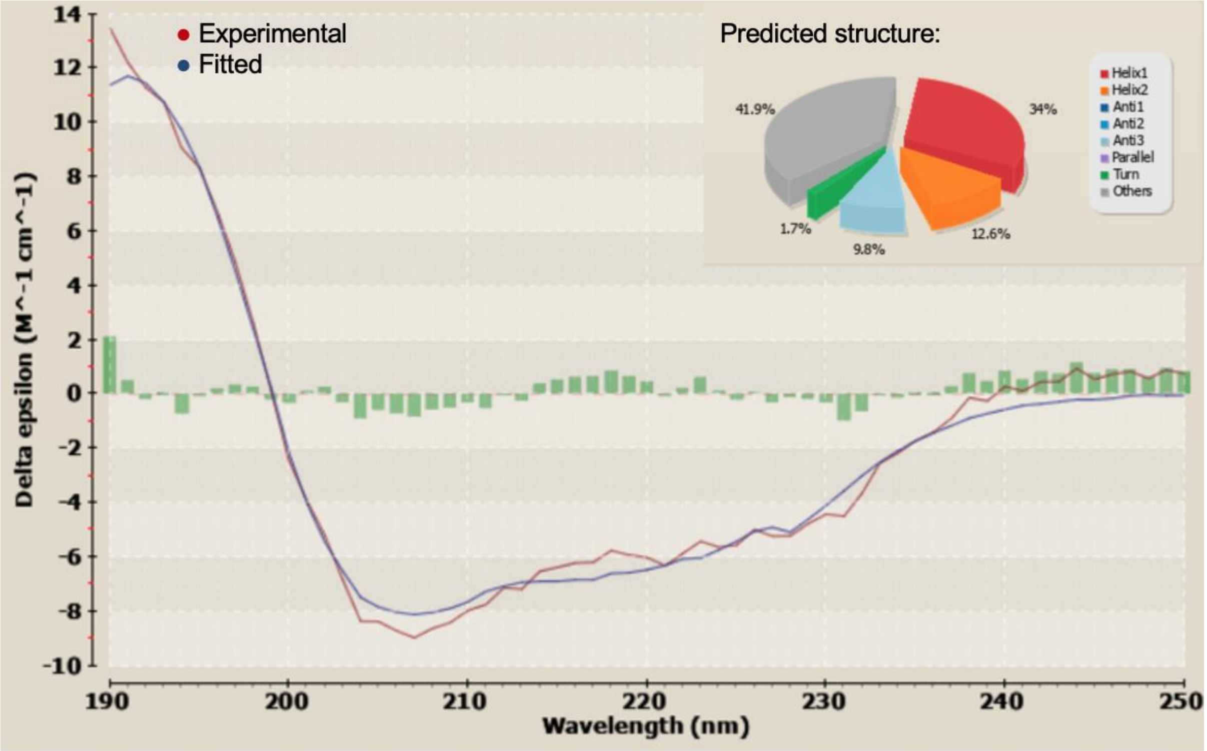


**Fig. S7** Alternate analysis of circular dichroism data. The circular dichroism data shown in fig 9 for Fny-CMV 2b protein in the absence of TFE was also run through the BeStSel program (v1.3.230210). This program predicted a similar level of disorder within the 2b sequence (41.9 %) but a larger percentage of alpha helical residues (46.6 %) and a smaller percentage of beta sheet (9.8 %). This program does not use datasets specifically trained on disordered sequences but the results do align more closely with the structural data obtained from NMR analysis.

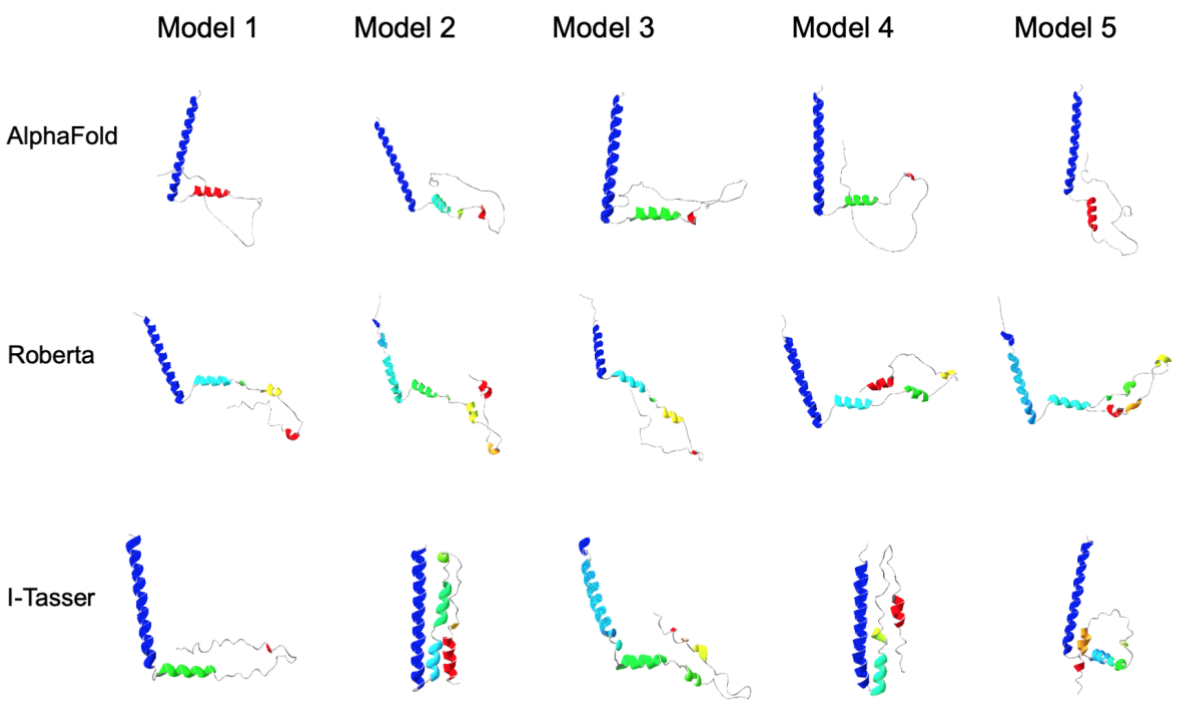


**Fig. S8.** Polypeptide folding predictions for the Fny-CMV 2b protein. Modelling predictions are shown from Alpha-Fold 2, RoseTTA fold and I-Tasser. All models show two alpha helixes in the N-terminal half of the 2b protein and the majority of models have a largely disordered C-terminal half.

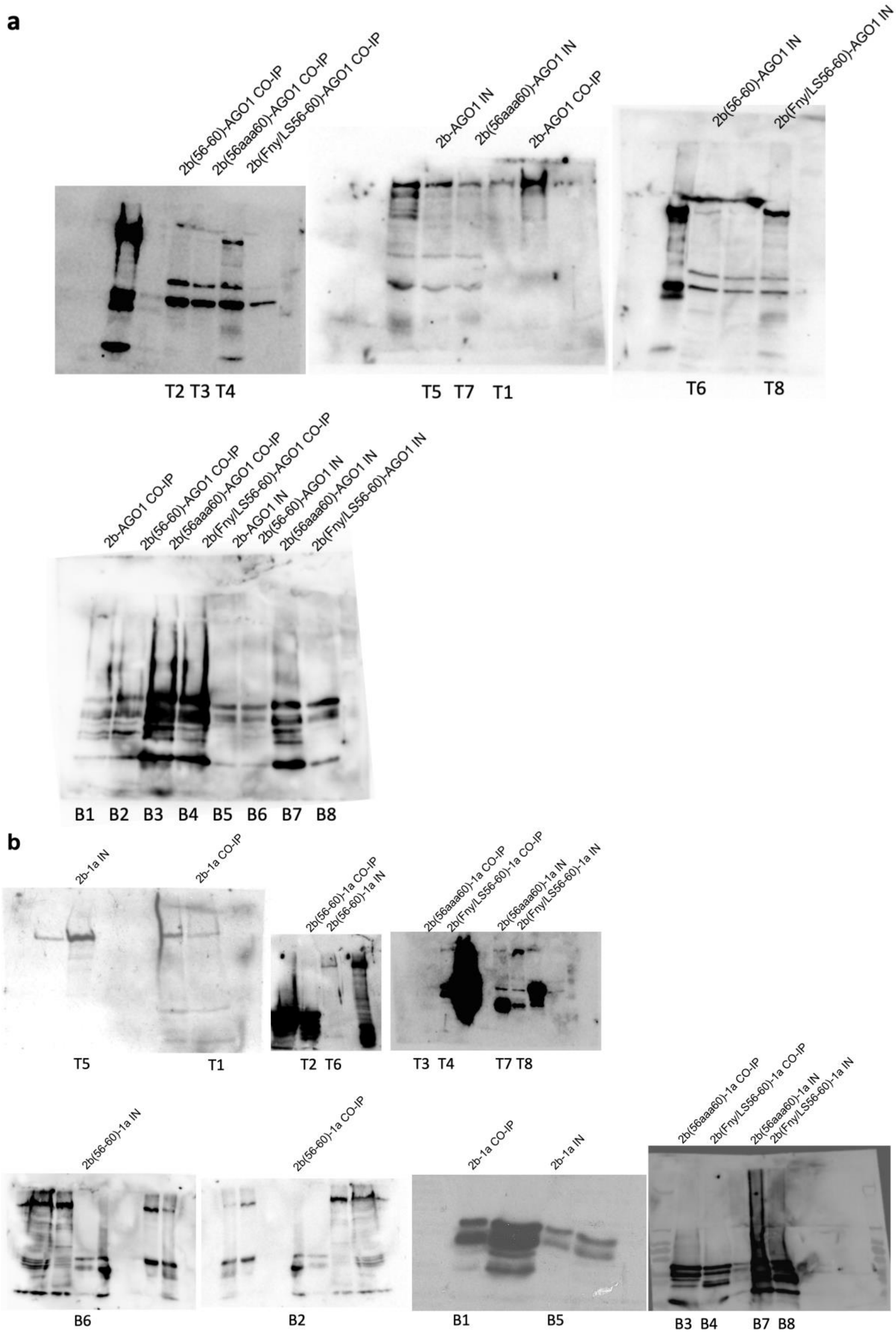
a
2b(56-60)-AGO1 CO-IP
2b(56aaa60)-AGO1 CO-IP
2b(Fny/LS56-60)-AGO1 CO-IP
T2 T3 T4
2b-AGO1 IN
2b(56aaa60)-AGO1 IN
2b-AGO1 CO-IP
T5 T7 T1
2b(56-60)-AGO1 IN
2b(Fny/LS56-60)-AGO1 IN
T6 T8
2b-AGO1 CO-IP
2b(56-60)-AGO1 CO-IP
2b(56aaa60)-AGO1 CO-IP
2b(Fny/LS56-60)-AGO1 CO-IP
2b-AGO1 IN
2b(56-60)-AGO1 IN
2b(56aaa60)-AGO1 IN
2b(Fny/LS56-60)-AGO1 IN
B1 B2 B3 B4 B5 B6 B7 B8
b
2b-1a IN
2b-1a CO-IP
T5
T1
2b(56-60)-1a CO-IP
2b(56-60)-1a IN
T2 T6
2b(56aaa60)-1a CO-IP
2b(Fny/LS56-60)-1a CO-IP
2b(56aaa60)-1a IN
2b(Fny/LS56-60)-1a IN
T3 T4
T7 T8
2b(56-60)-1a IN
B6
2b(56-60)-1a CO-IP
B2
2b-1a CO-IP
2b-1a IN
B1
B5
2b(56aaa60)-1a CO-IP
2b(Fny/LS56-60)-1a CO-IP
2b(56aaa60)-1a IN
2b(Fny/LS56-60)-1a IN
B3 B4
B7 B8

**Fig. S9** Whole gel images showing association of wild type and mutant Fny-CMV 2b proteins with Fny-CMV 1a or AGO1 proteins demonstrated by co-immunoprecipitation. **a.** RFP- tagged 2b proteins were co-expressed with GFP tagged AGO1 proteins in *N. benthamiana* leaves. Total protein was subjected to immunoprecipitation with GFP-Trap or RFP-Trap beads followed by immunoblot analysis with anti-RFP antibodies to detect 2b- RFP or anti-GFP antibodies to detect GFP-AGO1. Bands are labelled with the contents of the total protein extract and treatment: input sample (IN), Co-immunoprecipitation with interacting partner (CO-IP). The position where each band appears in the composite blot figures is denoted by T or B referring to the top or bottom row of gel strips. **b.** RFP- tagged 2b proteins were co-expressed with GFP tagged 1a proteins in *N. benthamiana* leaves. Total protein was subjected to immunoprecipitation with GFP-Trap or RFP-Trap beads followed by immunoblot analysis with anti-RFP antibodies to detect 2b-RFP or anti-GFP antibodies to detect GFP-1a. Bands are labelled with the contents of the total protein extract and treatment: input sample (IN) or Co-immunoprecipitation with interacting partner (CO-IP). The position where each band appears in the composite blot figures is denoted by T or B referring to the top or bottom row of gel strips.

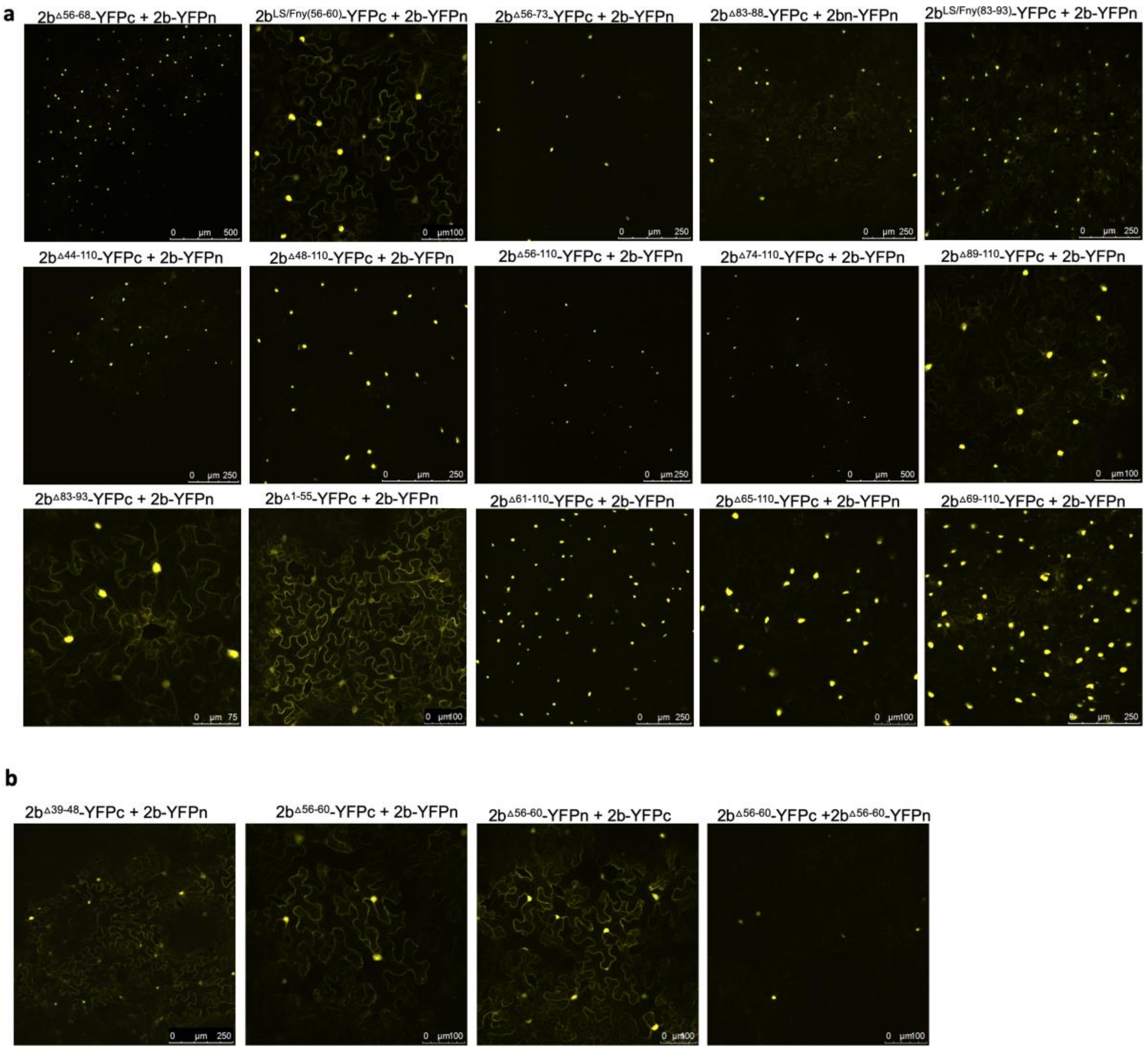


**Fig. S10** Mutations affecting the ability of 2b protein to interact with other CMV 2b proteins. Bimolecular fluorescence complementation was used to compare the self-interaction properties of the 2b proteins of Fny-CMV with mutant versions of the 2b protein using fusion proteins with the N- and C-terminal domains of the yellow fluorescent protein (2b-YFPn and 2b-YFPc). **a.** All mutant proteins were able to form heterodimers with WT 2b proteins in vivo, indicated by yellow fluorescence. The intracellular distributions of these homodimers were consistent with those seen for 2b-RFP or 2b-GFP mutant proteins and show a greater proportion of some mutant 2b proteins being present in the nucleus. **b.** Tagged mutant versions of the 2b protein lacking residues between 56-60 interacted strongly with tagged full length 2b proteins. However, self-interaction with two mutant versions of the 2b protein resulted in markedly reduced levels of self-interaction.

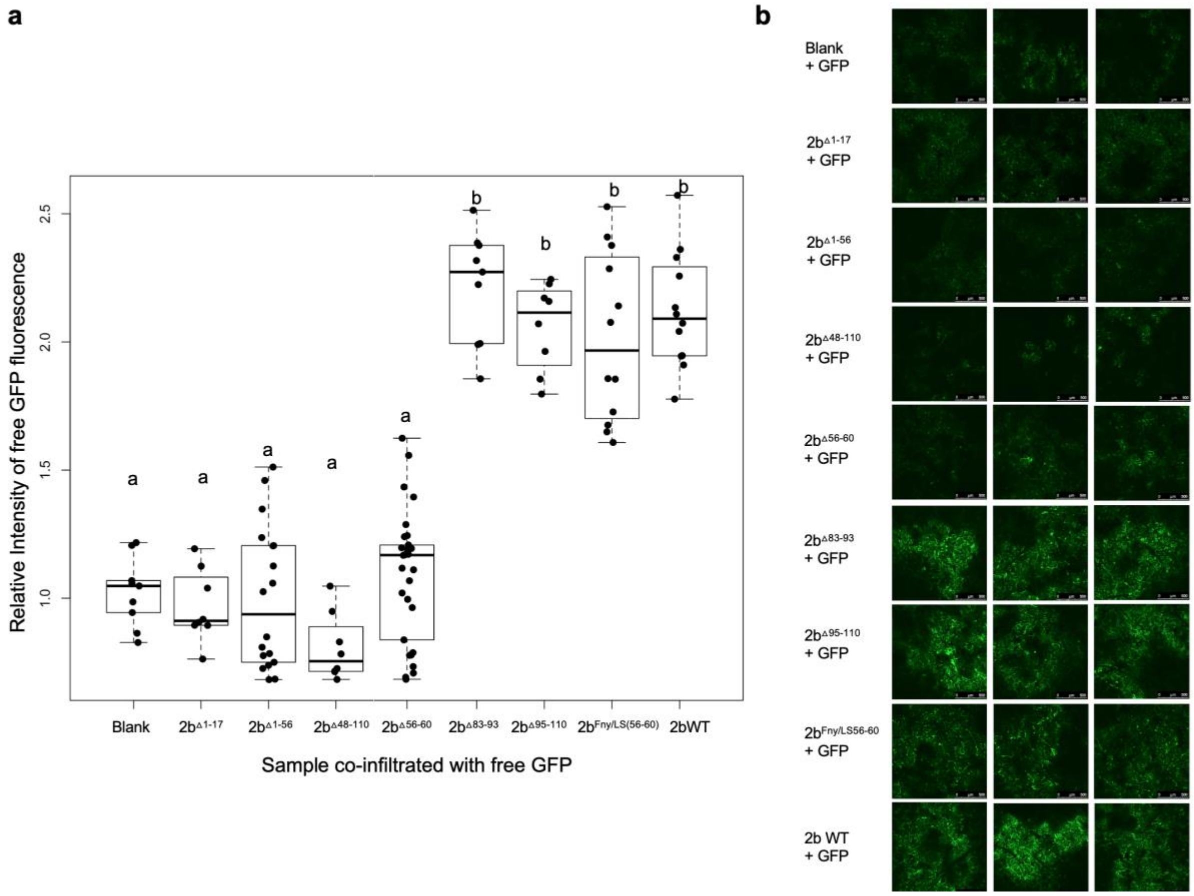


**Fig. S11** Mutations in the C-terminal region of the CMV 2b protein do not impact its RNA silencing suppressor activity. Green fluorescent protein (GFP) was expressed transiently, under a 35S promoter, in agroinfiltrated leaves of *Nicotiana benthamiana*. **a.** The relative intensity of GFP fluorescence was quantified using ImageJ as the integrated density (IntDen) of each image, for each treatment 12 days after infiltration. Individual relative fluorescence values are presented as jitter plots with each mean value and standard error depicted as black bars. Compared to the intensity of fluorescence emitted by leaves expressing GFP only, the relative intensity values of GFP fluorescence emitted by leaf tissue agroinfiltrated with mixtures of *A. tumefaciens* cells that included those carrying constructs expressing the full length 2b protein (WT), 95-110, 83-93 or Fny/LS(56-60) mutant versions of the 2b protein were significantly greater. Lower case letters *a* and *b* indicate mean values for relative fluorescence intensity that are significantly different from each ($P < 0.0001$: Tukey's multiple comparison of means). Values labelled with the same letter are not significantly different from each other. Number of independent leaves imaged for each treatment, $n > 6$. **b.** Typical confocal images of GFP fluorescence in the presence of full length or mutant versions of the CMV 2b protein as indicated.
